# Borgs are giant extrachromosomal elements with the potential to augment methane oxidation

**DOI:** 10.1101/2021.07.10.451761

**Authors:** Basem Al-Shayeb, Marie C. Schoelmerich, Jacob West-Roberts, Luis E. Valentin-Alvarado, Rohan Sachdeva, Susan Mullen, Alexander Crits-Christoph, Michael J. Wilkins, Kenneth H. Williams, Jennifer A. Doudna, Jillian F. Banfield

## Abstract

Anaerobic methane oxidation exerts a key control on greenhouse gas emissions ^1^, yet factors that modulate the activity of microorganisms performing this function remain little explored. In studying groundwater, sediments, and wetland soil where methane production and oxidation occur, we discovered extraordinarily large, diverse DNA sequences that primarily encode hypothetical proteins. Four curated, complete genomes are linear, up to ~1 Mbp in length and share genome organization, including replicore structure, long inverted terminal repeats, and genome-wide unique perfect tandem direct repeats that are intergenic or generate amino acid repeats. We infer that these are a new type of archaeal extrachromosomal element with a distinct evolutionary origin. Gene sequence similarity, phylogeny, and local divergence of sequence composition indicate that many of their genes were assimilated from methane-oxidizing *Methanoperedens* archaea. We refer to these elements as “Borgs”. We identified at least 19 different Borg types coexisting with *Methanoperedens* in four distinct ecosystems. Borg genes expand redox and respiratory capacity (e.g., clusters of multiheme cytochromes), ability to respond to changing environmental conditions, and likely augment *Methanoperedens* capacity for methane oxidation (e.g., methyl coenzyme M reductase). By this process, Borgs could play a previously unrecognized role in controlling greenhouse gas emissions.

## Introduction

Of all of Earth’s biogeochemical cycles, the methane cycle may be most tightly linked to climate. Methane (CH_4_) is a greenhouse gas roughly 30 times more potent than carbon dioxide (CO_2_), and approximately 1 gigaton is produced annually by methanogenic (methane-producing) archaea that inhabit anoxic environments^2^. The efflux of methane into the atmosphere is mitigated by methane-oxidizing microorganisms (methanotrophs). In oxic environments CH_4_ is consumed by aerobic bacteria that use a methane monooxygenase (MMO) and O_2_ as terminal electron acceptor ^3^, whereas in anoxic environments <u>an</u>aerobic <u>me</u>thanotrophic archaea (ANME) use a reverse methanogenesis pathway to oxidize CH_4_, the key enzyme of which is methyl-CoM reductase (MCR) ^4,5^. Some ANMEs rely on a syntrophic partner to couple CH_4_ oxidation to the reduction of terminal electron acceptors, yet *Methanoperedens* (ANME-2d, phylum *Euryarchaeota*) can directly couple CH_4_ oxidation to the reduction of iron, nitrate or manganese ^6,7^. Some phenomena have been suggested to modulate methane oxidation rates. For example, some phages can decrease methane oxidation rates by infection and lysis of methane-oxidizing bacteria ^8^ and others with the critical subunit of MMO ^9^ likely increase the ability of their host bacteria to generate energy during phage replication. Here, we report the discovery of novel extrachromosomal elements that clearly replicate within *Methanoperedens*. Their numerous and diverse metabolism-relevant genes, huge size, and distinctive genome architecture distinguish these archaeal extrachromosomal elements from all previously reported elements associated with archaea ^10–12^ and from bacteriophages, which typically have one or a few biogeochemically relevant genes ^13,14^. We infer that these novel extrachromosomal elements likely augment *Methanoperedens* capacities, and thus increase methane oxidation rates.

### Genome Structure and Features

By analysis of whole-community metagenomic data from saturated vernal pool (wetland) soils in CA (**Fig. S1**), we discovered enigmatic genetic elements, the genomes for three of which were carefully manually curated to completion (**methods**). From sediment samples from the Rifle, CO aquifer ^15^, we recovered partial genomes from a single population related to those from the vernal pool soils; the sequences were combined and manually curated to ultimately yield a fourth complete genome (**methods**). All four curated genomes are linear and terminated by >1 kbp inverted repeats. The genome sizes range from 661,708 to 918,293 kbp (**Fig. 1A; Table 1; Table S1**). Prominent features of all genomes are tens of regions composed of perfect tandem direct repeats (**Fig. 1B; Table S2)**that are overwhelmingly novel (**Fig. S2**) and occur in both intergenic regions and in genes where they usually introduce perfect amino acid repeats (**Table S2**). All genomes have two replicores of unequal lengths and initiate replication at the chromosome ends (**Fig S3**). Each replicore carries essentially all genes on one strand (**Fig. 1A)**. Although the majority of genes are novel, ~21% of the predicted proteins have best matches to proteins of Archaea (**Fig. S4A**), and the vast majority of these have best matches to proteins of *Methanoperedens* (**Fig. S4B**). Notably, the GC contents of the four genomes are ~10% lower than those of previously reported and coexisting *Methanoperedens* species (**Fig. 2A**). The sequences are much more abundant in deep, anaerobic soil samples (**Fig. S5)**and show no consistent relationship to abundances of coexisting *Methanoperedens* (**Fig. 2B**). Individually, their abundance can exceed that of the most abundant *Methanoperedens* by up to ~8X. We rule out the possibility that these sequences represent genomes of novel Archaea, as they lack almost all of the single copy genes found in archaeal genomes and sets of ribosomal proteins that are present even in obligate symbionts **(Figs. S6, S7A, Tables S3-S6**). There are no additional sequences in the datasets that could comprise additional portions of these genomes. Thus, they are clearly neither part of *Methanoperedens* genomes nor parts of the genomes of other archaea.

**Figure 1:**
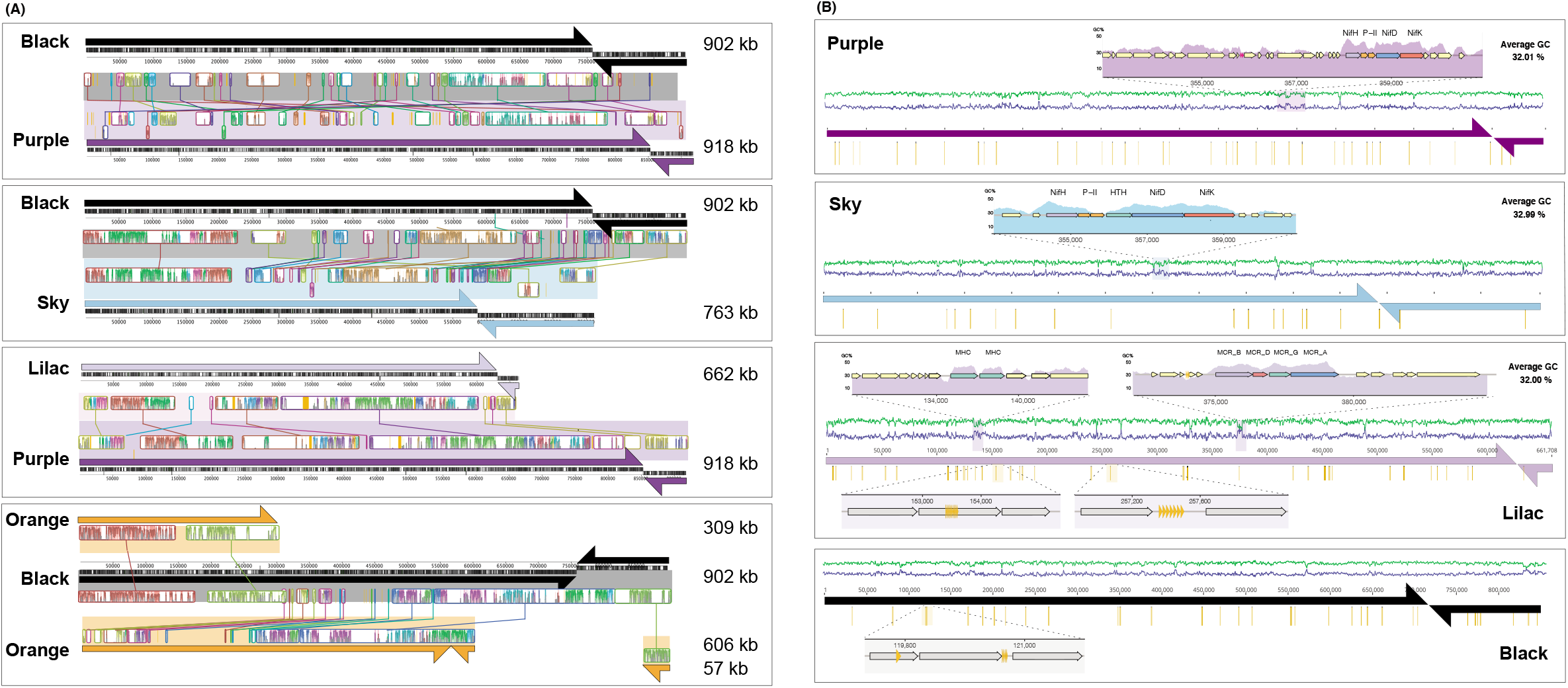
Borgs share overall genomic features. (A) Genome replicores (arrows) and coding strands (black bars) for aligned pairs of the four complete (Black, Purple, Sky and Lilac) and one near-complete (Orange) Borg. Blocks of sequence with identifiable similarity are shown in between each pair (colored graphs linked by lines, y-axes show similarity). (B) Genome overviews showing the distribution of three or more perfect tandem direct repeats (gold rods) along the complete genomes. Insets provide examples of local elevated GC content associated with certain gene clusters and within gene and intergenic tandem direct repeats (gold arrows).

**Figure 2:**
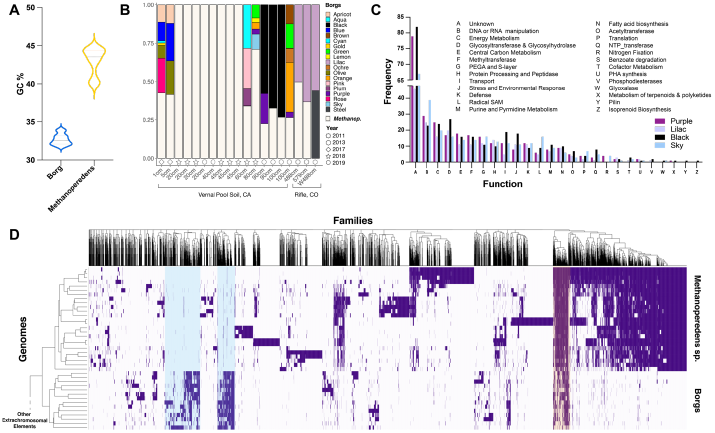
Borg and *Methanoperedens* genomic features and abundance patterns. A. The average genome GC contents of Borgs and *Methanoperedens* are distinct. B. Relative abundances of *Methanoperedens* and Borgs in samples collected over time and arrayed by sample collection depth from the vernal pool soils, sediments and groundwater. The absolute abundances of Borgs are far greater in the deeper compared to shallower soils (**Figure S5**). C. Frequency of genes in different functional groups in the four complete Borg genomes. D. Comparison of the protein family composition of Borgs and *Methanoperedens*. Clustering based on shared protein family content highlights groups of Borg-specific protein families (blue shading) and protein families shared with their hosts (orange shading). The full clustering, including diverse archaeal mobile elements, is shown in **Figure S6.**

**Table 1:**
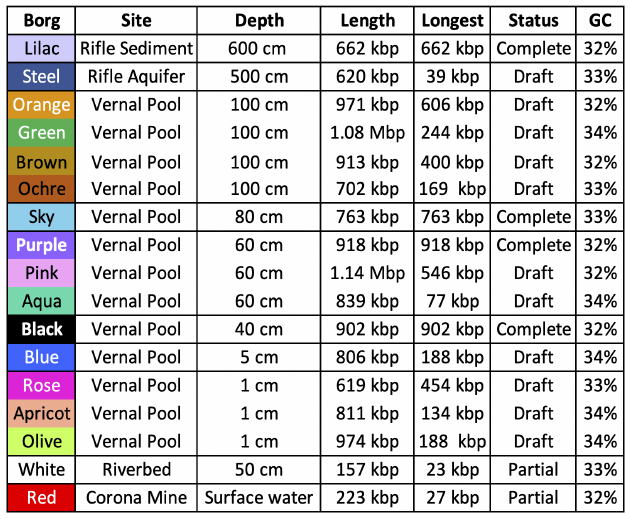
Manually curated complete and draft genomes for the best sampled Borgs. Length is the genome length. Longest is the size of the largest genome fragment. Status indicates degree of genome completeness: complete genomes have been corrected and fully verified throughout. GC is the genome-wide average GC content. For details for these and less abundant examples, see **Table S1**.

| **Borg** | **Site** | **Depth** | **Length** | **Longest** | **Status** | **GC** |
| --- | --- | --- | --- | --- | --- | --- |
| Lilac | Rifle Sediment | 600 cm | 662 kbp | 662 kbp | Complete | 32% |
| Steel | Rifle Aquifer | 500 cm | 620 kbp | 39 kbp | Draft | 33% |
| Orange | Vernal Pool | 100 cm | 971 kbp | 606 kbp | Draft | 32% |
| Green | Vernal Pool | 100 cm | 1.08 Mbp | 244 kbp | Draft | 34% |
| Brown | Vernal Pool | 100 cm | 913 kbp | 400 kbp | Draft | 32% |
| Ochre | Vernal Pool | 100 cm | 702 kbp | 169 kbp | Draft | 33% |
| Sky | Vernal Pool | 80 cm | 763 kbp | 763 kbp | Complete | 33% |
| Purple | Vernal Pool | 60 cm | 918 kbp | 918 kbp | Complete | 32% |
| Pink | Vernal Pool | 60 cm | 1.14 Mbp | 546 kbp | Draft | 32% |
| Aqua | Vernal Pool | 60 cm | 839 kbp | 77 kbp | Draft | 34% |
| Black | Vernal Pool | 40 cm | 902 kbp | 902 kbp | Complete | 32% |
| Blue | Vernal Pool | 5 cm | 806 kbp | 188 kbp | Draft | 34% |
| Rose | Vernal Pool | 1 cm | 619 kbp | 454 kbp | Draft | 33% |
| Apricot | Vernal Pool | 1 cm | 811 kbp | 134 kbp | Draft | 34% |
| Olive | Vernal Pool | 1 cm | 974 kbp | 188 kbp | Draft | 34% |
| White | Riverbed | 50 cm | 157 kbp | 23 kbp | Partial | 33% |
| Red | Corona Mine | Surface water | 223 kbp | 27 kbp | Partial | 32% |

A few percent of the genes in the genomes have locally elevated GC contents that approach, and in some cases match, those of coexisting *Methanoperedens* (**Fig. 1B**). This, and very high similarity of some protein sequences to those of *Methanoperedens*, indicates that these genes were acquired by lateral gene transfer from *Methanoperedens*. Other genes with best matches to *Methanoperedens* genes have lower GC contents (closer to those of Borgs at ~33%), suggesting that their DNA composition has partly or completely ameliorated since acquisition ^16^.

Archaeal extrachromosomal elements include viruses ^17^, plasmids ^18^ and minichromosomes, sometimes also referred to as megaplasmids ^10–12^. The genomes reported here are much larger than those of all known archaeal viruses, some of which have small, linear genomes ^12^, and at least three are larger than any known bacteriophage ^19^. These linear elements are larger than all of the reported circular plasmids that affiliate with halophiles, methanogens and thermophiles. They lack distinguishable viral or plasmid proteins, replication origins, rRNA loci, and essential genes typical of archaeal minichromosomes ^12^. Moreover, the protein family profiles are quite distinct from those of archaeal extrachromosomal elements (**Fig. 2D, Fig. S6**). Some bacterial megaplasmids have been reported to be large and linear and can contain repeats ^20^, but the repeats are interspaced (i.e., not tandem) and they typically do not encode essential genes ^21^. Given all of these observations, we conclude that the genomes represent novel archaeal extrachromosomal elements that occur in association with, but not as part of, *Methanoperedens* genomes. We refer to these as Borgs, a name that reflects their propensity to assimilate genes from organisms, most notably *Methanoperedens*.

Using criteria based on the features of the four complete Borgs, we searched for additional Borgs in our metagenomic datasets from a wide diversity of environment types. From the vernal pool soil, we constructed bins for 11 additional Borgs, some of which exceed 1 Mbp in length (**Table 1, Table S1**). Other Borgs were sampled from the Rifle, CO aquifer, discharge from an abandoned Corona mercury mine in Napa County, CA, and from shallow riverbed pore fluids in the East River, CO. In total, we recovered genome bins for 19 different Borgs, each of which was assigned a color-based name. Interestingly, we found no Borgs in some samples, despite the presence of *Methanoperedens* at very high abundance levels (**Fig. S7B**). Thus, it appears that these extrachromosomal elements do not associate with a monophyletic group of *Methanoperedens*.

Pairs of the four complete Borg genomes (Purple, Black, Sky and Lilac) and three fragments of the Orange Borg are alignable over much of their lengths (**Fig 1A**). Intriguingly, despite only sharing <50% average nucleotide identity across most of their genomes, the curated Rose and fully curated Sky Borgs have multiple regions that share 100% nucleotide identity, one of which is ~11 kbp in length. This suggests that these two Borgs recombined, indicating that they recently co-existed within the same host cell (**Fig S8A,B**).

### Borg gene inventories

Many Borg genomes encode mobile element defense systems, including RNA-targeting type III-A CRISPR-Cas systems that lack spacer acquisition machinery, a feature previously noted in huge bacterial viruses^19^. An Orange Borg CRISPR spacer targets a gene in a mobile region in a coexisting *Methanoperedens* (**Fig. S8C**), further supporting the conclusion that *Methanoperedens* are the Borg hosts.

The four complete genomes and almost all of the near-complete and partial genomes encode ribosomal protein L11 (rpL11), and some have one or two other ribosomal proteins **(Fig. S7A)**. The rpL11 protein sequences form a group that places phylogenetically sibling to those of *Methanoperedens* (**Fig. S9**), further reinforcing the link between Borgs and *Methanoperedens*. Four additional rpL11 sequences identified on short contigs from the vernal pool group with the Borg sequences and likely represent additional Borgs (**Table S1**). The topology of the rpL11 tree, and similar topologies observed for phylogenetic trees constructed using other ribosomal proteins and translation-related genes, may indicate the presence of translation-related genes in the Borg ancestor **(Fig. S7A; Fig. S9)**.

The most highly represented Borg genes are glycosyltransferases, genes involved in DNA and RNA manipulation, transport, energy, and the cell surface (PEGA and S-layer proteins). Also prevalent are many membrane-associated proteins of unknown function that may impact the membrane profile of their host (**Fig. 2C**). At least seven Borgs carry a *nifHDK* operon for nitrogen fixation, also predicted in *Methanoperedens* genomes (**Fig. 1B, Fig. S10, Table S6**). Potentially related to survival under resource limitation are genes in at least 10 Borg genomes for synthesis of the carbon storage compound polyhydroxyalkanoate, a capacity also predicted for *Methanoperedens*^22^. Other stress-related genes encode tellurium resistance proteins that do not occur in *Methanoperedens* genomes (**Table S5**). Intriguingly, all Borgs carry large FtsZ-tubulin homologs that may be involved in membrane remodeling or division, and proteins that resemble Major Vault Proteins and the TEP1-like TROVE domain protein that also do not occur in *Methanoperedens* genomes (**Table S5**). These are known to form the highly conserved and enigmatic eukaryotic vault organelle, a ribonucleoprotein that has been suggested to be involved in multidrug resistance, nucleo-cytoplasmic transport, mRNA localization, and innate immunity ^23^. Several Borgs encode two genes of the TCA cycle (citrate synthase and aconitase, **Fig. S10C)**.

Many Borg genes are predicted to play roles in redox and respiratory reactions. The Black Borg encodes *cfbB* and *cfbC*, genes involved in biosynthesis of F430, the cofactor for methylcoenzyme M reductase (MCR), the central enzyme involved in methane oxidation by *Methanoperedens*. The similarity in GC content of Borg *cfbB* and *cfbC* and protein sequences of coexisting *Methanoperedens* suggests that these genes were acquired from *Methanoperedens* recently. The Blue and Olive Borgs encode *cofE* (coenzyme F420:L-glutamate ligase), which is involved in biosynthesis of a precursor for F430. The Blue and Pink Borgs have an electron bifurcating complex **(Fig. S10B)**that includes D-Lactate dehydrogenase. Eight Borgs encode genes for biosynthesis of tetrahydromethanopterin, a coenzyme used in methanogenesis, and ferredoxin proteins which may serve as electron carriers. The Green and Sky Borgs also encode 5,6,7,8-tetrahydromethanopterin hydro-lyase (Fae), an enzyme responsible for formaldehyde detoxification. Also identified were genes encoding carbon monoxide dehydrogenase (CODH), plastocyanin, cupredoxins and many multiheme cytochromes (MHC). These results indicate substantial Borg potential to augment the energy generation by *Methanoperedens*. This is especially apparent for the Lilac Borg.

### Lilac Borgs potential to augment *Methanoperedens* function

We analyzed the genes of the complete Lilac Borg genome in detail as, unlike the other Borgs, the Lilac Borg co-occurs with a single group of relatively abundant *Methanoperedens* that likely represent the host (**Fig. 3, Table S7**). Remarkably, this Borg genome encodes an MCR complex, which is central to methanogenesis and reverse methanogenesis. The *mcrBDGA* cluster shares high (75-88%) amino acid sequence identity with that of the coexisting *Methanoperedens* genome. This complex is also encoded by a fragment of the Steel Borg. For both the Lilac and Steel Borgs, the GC content of the region encoding this operon is elevated relative to the average Borg values. *Methanoperedens* pass electrons from methane oxidation to terminal electron acceptors (Fe^3+^, NO_3_^-^ or Mn^4+^) via MHC ^24–26^. The Lilac Borg genome encodes 16 multiheme cytochromes with up to 32 heme-binding motifs within one protein. Membrane-bound and extracellular MHC likely diversify the range of *Methanoperedens* extracellular electron acceptors, potentially increasing the rate of methane oxidation.

**Figure 3:**
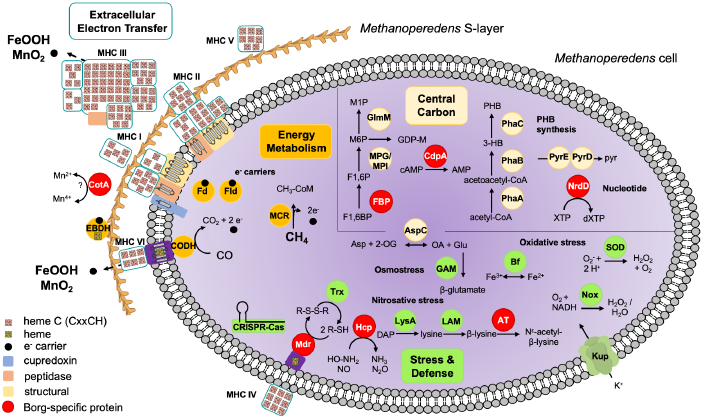
Cell cartoon illustrating capacities inferred to be provided to *Methanoperedens* by coexisting Lilac Borgs. Like all Borgs, this Borg lacks the capacity for independent existence, and we infer that it exists within host *Methanoperedens* cells. Borg specific proteins (red circles) are those that were not identified in the genome of coexisting *Methanoperedens*. Borg-encoded capacities are grouped into the major categories of energy metabolism (including the MCR complex involved in methane oxidation), extracellular electron transfer (including multiheme cytochromes, MHC) involved in electron transport to external electron acceptors), central carbon metabolism (including the ability to make polyhydroxybutyrate, PHB), and stress response/defense (including production of compatible solutes). Locus codes are listed in **Table S7**.

The Lilac Borg encodes a functional NiFe CODH, but this is fragmented in some genomes. Other genes for the acetyl-CoA decarbonylase/synthase complex are present only in *Methanoperedens*. The CODH is located in proximity to a cytochrome *b* and cytochrome *c*, so electrons from CO oxidation could be passed to an extracellular terminal acceptor such as Fe^3+^in an energetically downhill reaction. This would allow removal of toxic CO and may contribute to the formation of a proton gradient that can be harnessed for energy conservation.

The Lilac Borg has a gene resembling the gamma subunit of ethylbenzene dehydrogenase (EBDH), which is involved in transferring electrons liberated from the hydroxylation of ethylbenzene and propylbenzene ^27^. This EBDH-like protein is located extracellularly, and given heme binding and cohesin domains, it may be involved in electron transfer and attachment.

Although the Lilac Borg lacks genes for a nitrate reductase, it encodes a probable hydroxylamine reductase (Hcp) that may scavenge toxic NO and hydroxylamine byproducts of *Methanoperedens* nitrate metabolism. As the *hcp* gene was not identified in coexisting *Methanoperedens*, the Borg gene may protect *Methanoperedens* from nitrosative stress. Proteins such as H_2_O_2_-forming NADH oxidase (Nox) and superoxide dismutase (SOD) may protect against reactive oxygen species. An alkylhydroperoxidase, two probable disulfide reductases, and a bacterioferritin all may detoxify the H_2_O_2_ byproduct of Nox and SOD. The Lilac Borg also encodes genes that likely augment osmotic stress tolerance. This Borg, but not *Methanoperedens*, has the ability to make N^ε^-acetyl-β-lysine as an osmolyte. An aspartate aminotransferase links the tricarboxylic acid cycle and amino acid synthesis, producing glutamate that can be used for the production of the osmolyte β-glutamate. More importantly, perhaps, it has recently been established that this single enzyme can produce methane from methylamine ^28^, raising the possibility of methane cycling within the Borg - *Methanoperedens* system.

The Lilac Borg has three large clusters of genes. The first may be involved in cell wall modification, as it encodes large membrane-integral proteins with up to 17 transmembrane domains, proteins for polysaccharide synthesis, glycosyltransferases, and likely carbohydrate active proteins. The second contains key metabolic valves that connect gluconeogenesis with mannose metabolism for the production of glycans. One gene, fructose 1,6-bisphosphatase (FBP), was not identified in the *Methanoperedens* genomes and may regulate carbon flow from gluconeogenesis to mannose metabolism. In between these clusters are 12 genes with PEGA domains with similarity to S-layer proteins. Cell surface proteins, along with these PEGA proteins, account for ~13% of all Lilac Borg genes. We conclude that functionalities related to cell wall architecture and modification are key to the impact of these extrachromosomal elements on their host, perhaps triggering cell wall modification for adaptation to changing environmental conditions (**Fig. 3**).

## Conclusions

Borgs are enigmatic extrachromosomal elements that can approach (and likely exceed) 1 Mbp in length (**Table 1**). We can neither prove that they are archaeal viruses or plasmids or mini-chromosomes, nor can we prove that they are not. Regardless of the name, they are clearly different from anything that has been reported previously. It is fascinating to ponder their possible evolutionary origins. Are they giant linear viruses or plasmids unlike anything previously reported? Alternatively, are they auxiliary chromosomes? Perhaps they were once a sibling *Methanoperedens* lineage that underwent gene loss and established a symbiotic association within *Methnoperedens*, as might be indicated by the monophyly of some of their *Methanoperedens*-like genes (**Figs. S9, S10**). We suspect that different Borgs tend to associate with different *Methanoperedens* species, yet Borg homologous recombination may indicate movement among hosts, thus their possible roles as gene transfer agents. It has been noted that *Methanoperedens* have been particularly open to gene acquisition from diverse bacteria and archaea ^6^, and Borgs may have contributed to this. The existence of Borgs encoding MCR demonstrates for the first time that MCR and MCR-like proteins for metabolism of methane and short-chain hydrocarbons can exist on extrachromosomal elements and thus could potentially be dispersed across lineages, as is inferred to have occurred several times over the course of archaeal evolution ^17,29^. Borgs carry numerous metabolic genes, some of which produce variants of *Methanoperedens* proteins that could have distinct biophysical and biochemical properties. Assuming that these genes extend and augment *Methanoperedens* energy metabolism, Borgs may have far-reaching biogeochemical consequences, including reducing methane fluxes, with important and unanticipated climate implications.

## Acknowledgements

We thank Gene Tyson, Dipti Nayak, Nitin Baliga, Jamie Cate, and Spencer Diamond for helpful discussion, Elliot Smith, who proposed the name “Borg”, and Lily Law and Shufei Lei for data management assistance. We thank Yuki Amano for permission to mention *Methanoperedens*-dominated metagenomic datasets in which we did not discover Borgs. This research was supported by an NSF Fellowship to BAS, DFG Fellowship to MCS, the US Department of Energy, Office of Science, Office of Biological and Environmental Research under Award Number DE-AC02-05CH11231, Chan Zuckerberg Biohub and Innovative Genome Institute, UC Berkeley.

## Author contributions

Metagenomic datasets were contributed by BAS, RS, LVA, ACC, JWR, MJW, SM, KHW, JFB. Genome binning was done by JFB, BAS, ACC. Genome curation was conducted by JFB and BAS and CRISPR-Cas analysis by BAS. Borg genome structure, taxonomic breakdowns, horizontal gene transfer, and general feature analyses were conducted by BAS and JFB. Phylogenetic analyses were conducted by BAS, and repeat sequence comparisons across Borgs were done by BAS and JFB. General Borg and *Methanoperedens* gene inventory and protein family analyses were done by BAS and JFB and Lilac Borg in-depth analysis was done by MCS. BAS, MCS and JFB wrote the manuscript, with input from all authors.

## Data Availability

Prior to publication, the genomes reported in this study can be accessed via https://ggkbase.berkeley.edu/BMp/organisms

## Materials and Methods

### Sampling and creation of metagenomic datasets

We analyzed sequences from sediments of an aquifer in Rifle, Colorado that were retrieved from cores from depths of 5 and 6 m below the surface ^15^ in July 2011, and cell concentrates from pumped groundwater from the same aquifer collected at a time of elevated O_2_ concentration in May 2013. Discharge from the Corona Mine, Napa County, California was sampled in December, 2019. Shallow pore water was collected from the riverbed at the East River, Crested Butte, Colorado sampled in August 2016. Soil was sampled from depth intervals between 1 cm to 1 m from a permanently moist and at times water filled vernal pool located in Lake County, California. Vernal pool (wetland) soils were sampled in late October and early November of 2017, 2018 and 2019. DNA was extracted from each sample and submitted for Illumina sequencing (150 bp or 250 bp reads) at the QB3 facility, University of California, Berkeley. Reads were adapter and quality trimmed using BBduk ^30^ and sickle ^31^. Filtered reads were assembled using IDBA-UD ^32^ and MEGAHIT, gene predictions were established using Prodigal ^33^ and USEARCH ^34^ was used for initial annotations^31,32,34,35^. Functional predictions and predictions of tRNAs followed previously reported methods^19^.

### Genome identification, binning, and curation

Hundreds of kbp *de novo* assembled sequences were identified to be of interest as potential novel extrachromosomal elements first based on their taxonomic profile. The taxonomic profiles were determined through a voting scheme in which the taxonomy is assigned at the species to domain level (Bacteria, Archaea, Eukaryotes, no Domain) by comparison with a sequence database (protein annotations in the UniProt and ggKbase: https://ggkbase.berkeley.edu/) when the same taxonomic assignment received >50% votes. Assembled sequences selected for further analysis had no taxonomic profile, even at the Domain level. The majority of contigs of interest had more genes with similarity to those of archaea of the genus *Methanoperedens* than to any other genus (see **Fig. S4**). The second feature of interest was dominance by hypothetical proteins yet absence of genes that would indicate identification as phage or viruses or plasmids.

These initially identified large fragments were manually curated to remove scaffolding gaps and local assembly errors, to extend and join contigs with the same profile, GC and coverage, and then to extend the near complete sequences fully into their long terminal repeats. The last step required reassignment of reads mapped at one end and at double depth to both ends. The fully extended sequences had no unplaced reads extending outwards, despite genome-wide deep coverage. Given this, and the absence of any fragments that could potentially be part of a larger genome, it was concluded that sequences represented linear genomes.

In more detail, our curation method involved mapping of reads to the *de novo* fragments and extension within gaps and at termini using previously unplaced reads that we added based on overlap or by relocation of misplaced reads (these could often be identified based on improper paired reads distances and/or wrong orientation). Local assembly errors were sought by visualization of the reads mapped throughout the assembly and identified based on imperfect read support, or where a subset of reads were partly discrepant and discrepancies involved sequences that were shared by tandem direct repeats of the same region (i.e., the tandem direct repeat regions were collapsed during assembly). *De novo* assembled sequences often ended in tandem direct repeat regions because repeats fragment assemblies. To resolve local assembly errors, gaps were inserted and reads relocated to generate the sequence required to fill the gaps. This ensured comprehensive essentially perfect agreement between reads and the final consensus sequence. In some cases, the tandem direct repeat regions had greater than expected depth of mapped reads and no reads spanned the flanking unique sequences. In these cases, the repeat number was approximated to achieve the expected read depth. GC skew, and cumulative GC skew were calculated for the fully manually curated complete genomes and the patterns were used to identify the origins and terminus of replication. The pattern of use of coding strands for genes (predicted in Bacterial Code 11) were compared to these origin and terminus predictions to resolve genome organization. The curated sequences were searched for perfect repeats of lengths ≥ 50 nucleotides using Repeat Finder in Geneious. When repeat sequences overlapped, the unit of direct repeat was identified and the length of that repeat, number of repeats, location (within gene vs. intergenic) and genome position were tabulated. Once the features characteristic of the extrachromosomal elements of interest had been determined, we sought related elements. Sequences of interest were identified based on (1) credible partial alignment with the complete sequences, (2) no Domain level profile, (3) GC content 30 - 35%, (4) regions with three or more direct tandem repeats scattered throughout the genome fragment and (5) more best hits to *Methanoperedens* proteins than to proteins from any other organisms. If scaffolds met criterion (1) they were immediately classified as targets. If they met most or all of the other criteria and had similar coverage values, they were binned together with other scaffolds from the same sample with these features. Often, ends of some of the contigs in the same bin overlapped perfectly and could be joined, increasing confidence in the bin quality. Genome sequences were aligned to each other using Mauve ^36^. Where anomalously high sequence identity suggestive of recent recombination was detected between Borgs, reads mapped to the region were visualized to verify that the assembly was correct (i.e., not chimeric). Genome fragments were phylogenetically profiled to establish relatedness to sequences in public databases. Sequences were classified as having no detectable hit if the protein had no similar database sequence with an e-value of <0.0001.

### CRISPR-Cas analysis

Borg and *Methanoperedens*-encoded CRISPR repeats and spacers were identified using CRISPRDetect ^37^. The coding sequences from this study were searched against Cas gene sequences reported from previous studies ^38^ using hmmsearch with *E* < 1 × 10-5 to identify the full locus. Matches were checked using a combination of hmmscan and BLAST searches against the NCBI nr database and manually verified by identifying colocated CRISPR arrays and Cas genes. Spacers extracted from between repeats of the CRISPR locus were compared to sequence assemblies from the sites where Borgs were identified using BLASTN-short ^39^. Matches with alignment length >24 bp and ≤1 mismatch were retained and targets were classified as bacteria, phage or other. CRISPR arrays that had ≤1 mismatch, were further searched for more spacer matches in the target sequence by finding more hits with ≤3 mismatches.

### Protein and gene content analysis

After the identification and curation of Borg genomes and accumulation of usearch annotations for coding sequences, functional annotations were further assigned by searching against PFAM r32, KEGG, pVOG. Transmembrane regions in proteins were predicted with TMHMM. All *Methanoperedens* genomes and genome assemblies as well as 1153 archaeal viruses and extrachromosomal elements were downloaded from the NCBI RefSeq database. Open reading frames were predicted using Prodigal, and all proteins from Borg genomes and the reconstructed ECE database were clustered into protein families and compared across genomes as previously described ^19^.

### Functional annotation

Genes of interest were further verified and compared using NCBI’s conserved domain search and InterproScan ^40^ to identify conserved motifs within the amino acid sequence. Multiheme cytochromes were identified based on ≥3 CxxCH motifs within one gene. The cellular localization of proteins was predicted with Psort (v3.0.3) using archaea as organism type. Proteins were compared using blastp and aligned using MAFFT v.7.407 ^41^ to visualize homologous regions and check conserved amino acid residues that constitute the active site or are required for cofactor/ligand binding.

### Phylogenetic trees

For each gene, references were compiled by BLASTing the corresponding gene against the NCBI nr database, and their top 50 hits clustered by CD-HIT using a 90% similarity threshold ^42^. The final set genes was aligned using MAFFT v.7.407 and a phylogenetic tree was inferred using IQTREE v.1.6.6 using automatic model selection ^43^

## EXTENDED DATA

**Figure S1.**
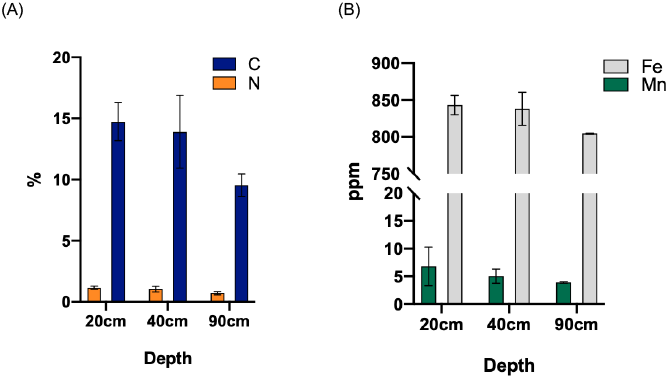
Geochemical profiles of the permanently moist and organic-rich vernal pool soils. (A) The concentrations of total carbon, nitrogen as well as (B) iron and manganese in vernal pool soils. Deeper soils, where these extrachromosomal elements are most abundant, are somewhat depleted in carbon, iron and manganese compared to shallow soils.

**Figure S2.**
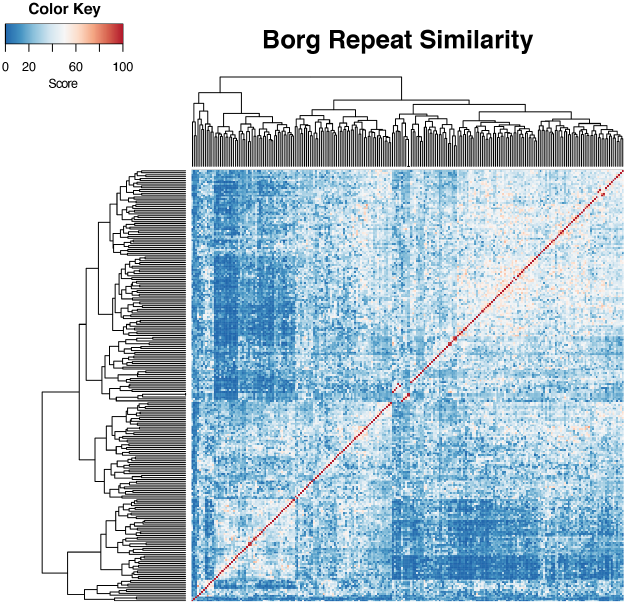
Sets of three or more perfect tandem direct repeats (TDR) are a characteristic feature of the Borg genomes. Up to 54 instances occur in the four complete Borg genomes, with, on average, one repeat every 12 (Lilac) - 31 (Sky) kbp. These repeat regions fragment assemblies and cause local assembly errors, which we resolved by manual curation (Methods). Within the TDR regions of the four curated, complete genomes, the unit repeats occur up to 20 times and unit repeats are up to 54 bps in length **(Table S2)**. The majority (54 - 64%) of these perfect TDRs are encoded in intergenic regions, although part or all of the first repeat may occur within the C-terminus of a protein-coding gene. When the TDRs occur within proteins, the unit lengths are almost always divisible by 3, so they introduce perfect amino acid repeats. TDR sequences within a single Borg genome are almost always unique. Repeat sequence comparison from the four complete curated Borgs highlights the novelty of almost all TDR sequences (both within and across genomes).

**Figure S3.**
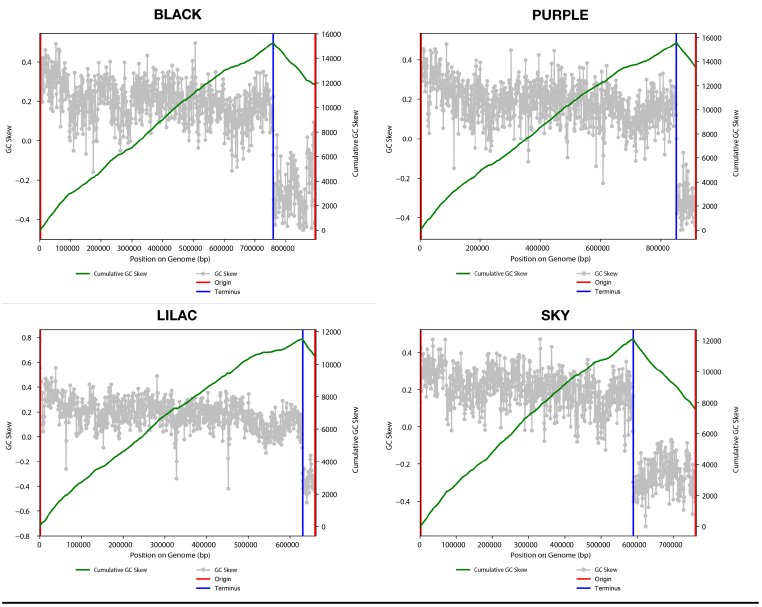
All genomes have two replicores of unequal lengths. GC skew (grey plots) and cumulative GC skew (green lines) across the four complete Borg genomes, all of which end in long inverted terminal repeats (1.4 - 2.7 kbp in length). The cumulative GC skew plots indicate replication is initiated in these terminal repeats (red lines). Blue lines mark the predicted replication termini. The red and blue lines define two replicores of unequal length that correspond almost completely to distinct coding strands (almost all genes on the +ve strand of the large replicore and on the −ve strand of the small replicore).

**Figure S4.**
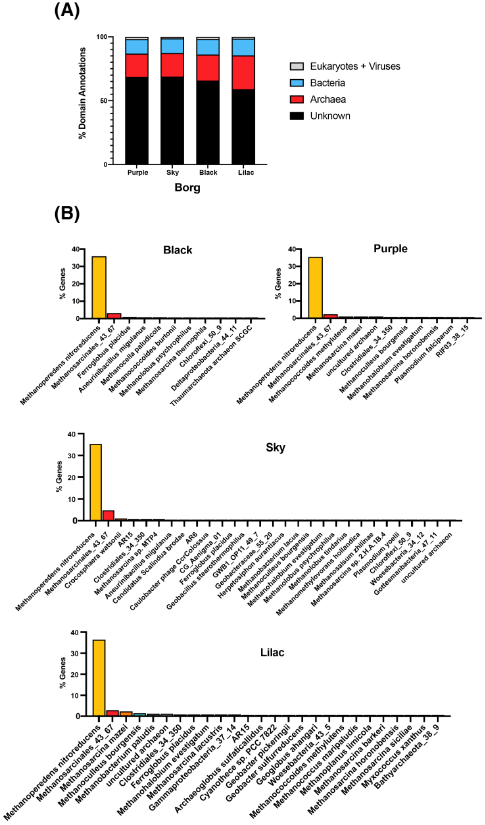
Taxonomic profiles of the four complete Borg genomes. **A.** In all cases, the majority of proteins have no similarity to proteins in the reference database (“Unknown”; e-value of >0.0001). For the cases where a protein has an identifiable hit (blue and red bars in A), the plots in **B** show the taxonomy of the organisms in which those hits were identified. Only cases where the same organism accounted for hits for >0.5% of genes are shown. The results clearly indicate that the vast majority of cases where proteins have identifiable matches involve matches to proteins of *Methanoperedens* (gold bars).

**Figure S5:**
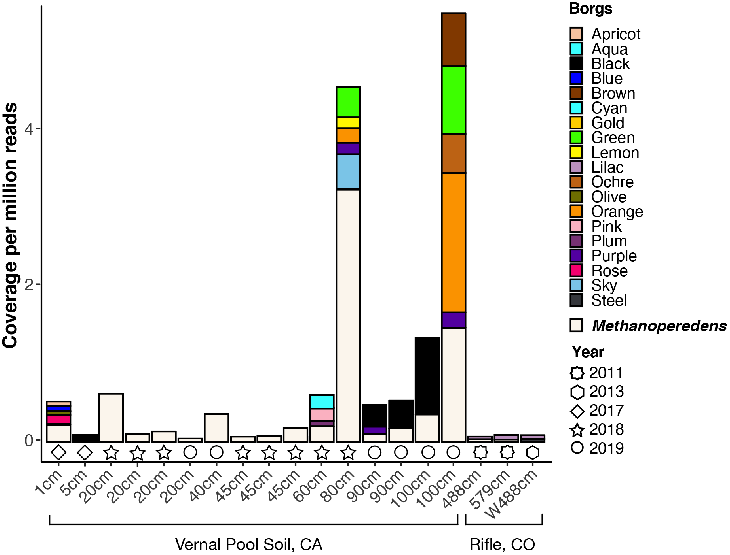
Abundance and distribution of Borgs and *Methanoperedens* in the vernal pool soil and Rifle aquifer. Note that the relative abundance of Borgs in soil samples is much higher in deeper soils. “W” indicates that the sample was pumped groundwater. Although some Borgs can substantially exceed all the combined abundance of *Methanoperedens*, no Borgs were detected in some *Methanoperedens*-bearing samples.

**Figure S6:**
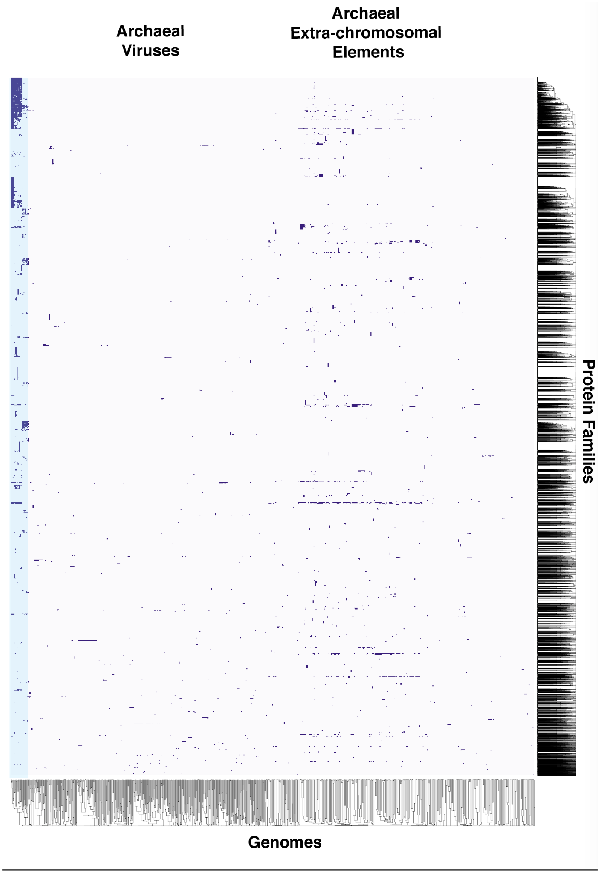
Comparison of the protein family content of *Methanoperedens*, Borgs, and other archaeal extrachromosomal elements. Shaded blocks indicate presence of the protein family in the corresponding genome. The blue highlight on the left side indicates that *Methanoperedens* (left) and Borg (right) protein family profiles are more similar to each other than, and distinct from, other extrachromosomal elements; for details see **Fig. 2D**.

**Figure S7:**
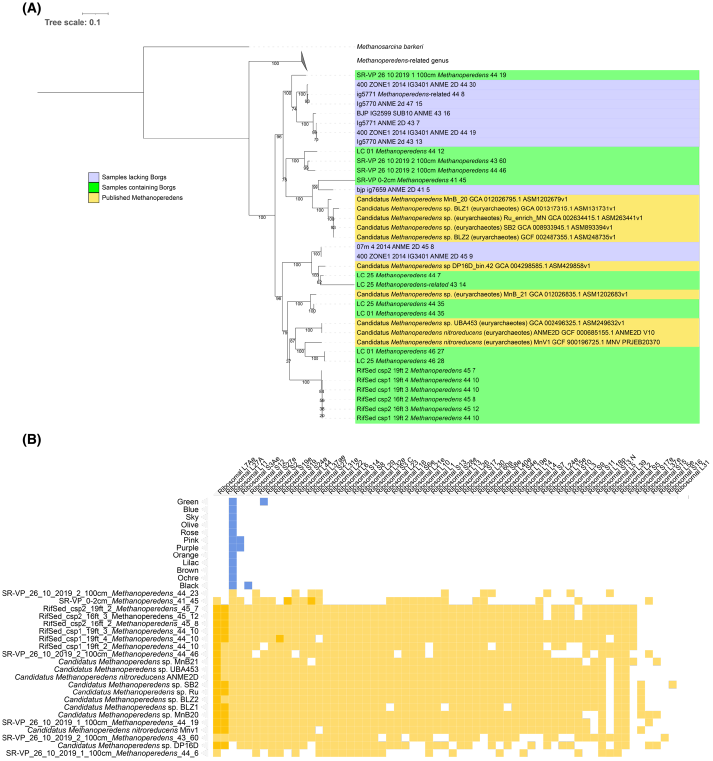
**(A)** The array of single-copy archaeal ribosomal genes (columns) vs. Borg (blue) and *Methanoperedens* (gold) genomes illustrating that although Borgs often have rpL11 and occasionally, other ribosomal proteins, they do not have the gene inventory needed to construct ribosomes. **(B)** Maximum Likelihood Phylogeny of concatenated ribosomal proteins from *Methanoperedens* species that do and do not coexist with Borgs and previously reported genomes. We found no data indicating the presence of Borgs in samples containing previously reported *Methanoperedens* genomes. We searched for Borgs in the samples highlighted in blue using the same methods used to detect Borgs in this study and concluded that they do not contain Borgs. A subset of the Borg-free samples contain *Methanoperedens* at very high abundance levels.

**Figure S8:**
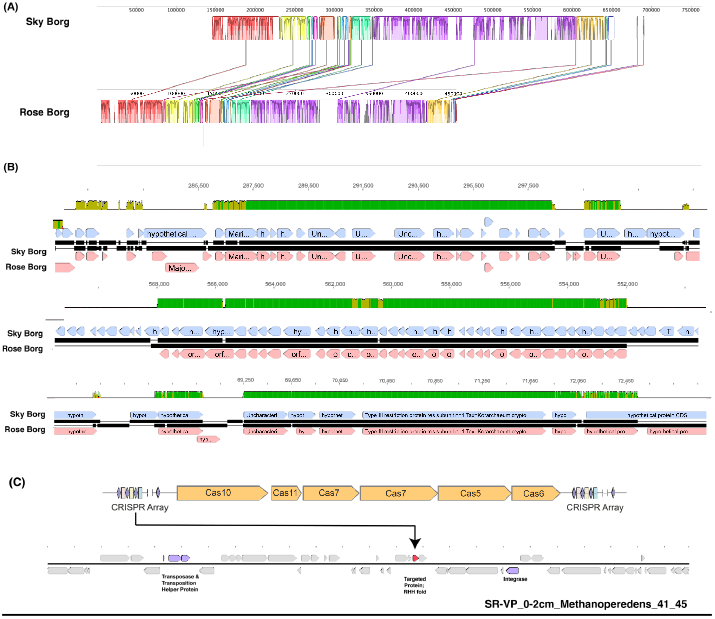
(A) Genome-to-genome comparisons provide evidence for recombination between two of the mostly closely related Borgs, Sky and Rose. These Borgs share only moderate overall genomic nucleic acid identity although, as is the case for other Borgs (**Figure 1A**), have blocks of partially alignable sequence throughout their genomes. Notable, and indicating recent homologous recombination, are 100% identical regions of up to ~11 kbp in length (B). Although not fully manually curated to completion, the relevant Rose Borg genome regions were carefully checked by inspection of the mapped reads to rule out chimeric assembly that could otherwise explain perfect identity with the Sky Borg sequence (Sky is one of the four curated complete genomes). (C) Diagram illustrating the organization of the Type III-A CRISPR-Cas system variant (lacking acquisition machinery and Csm6) in the Orange Borg. One spacer from the CRISPR array targets a small protein with a ribbon-helix-helix motif, a common transcriptional regulator in archaeal mobile elements, in a mobile region of a *Methanoperedens* genome bin from the same vernal pool site.

**Figure S9:**
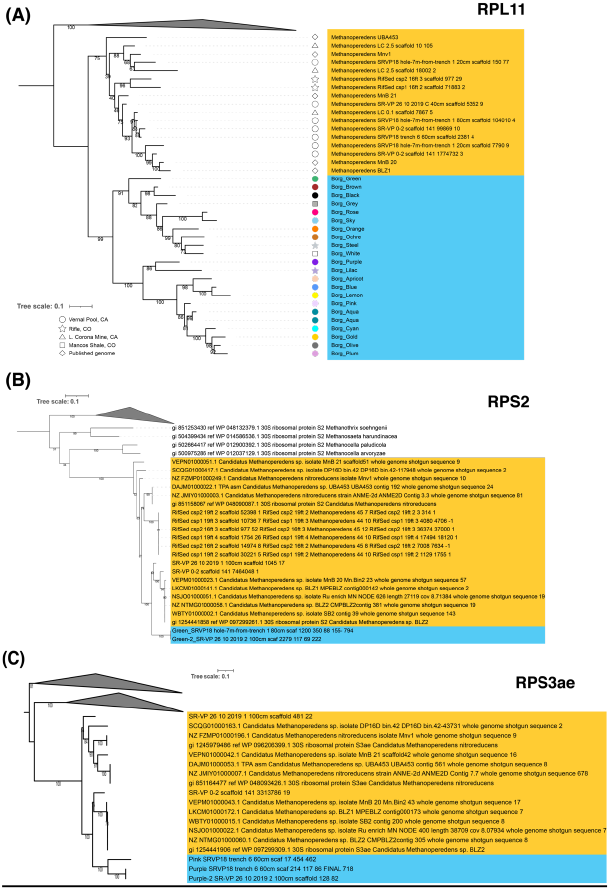
The Borg ribosomal sequences form monophyletic groups that cluster adjacent to those from *Methanoperedens*. Phylogenetic tree constructed using the protein sequences for (A) ribosomal protein L11 (rpL11), (B) Ribosomal protein S2 (C) Ribosomal protein 3ae.

**Figure S10:**
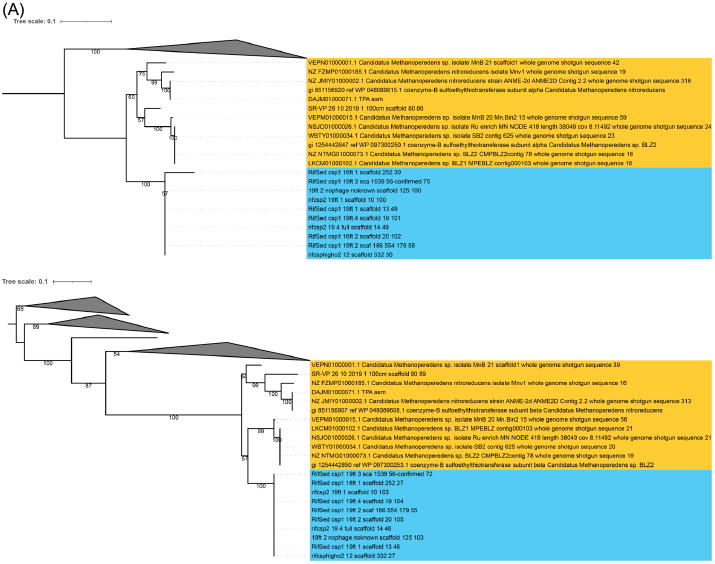
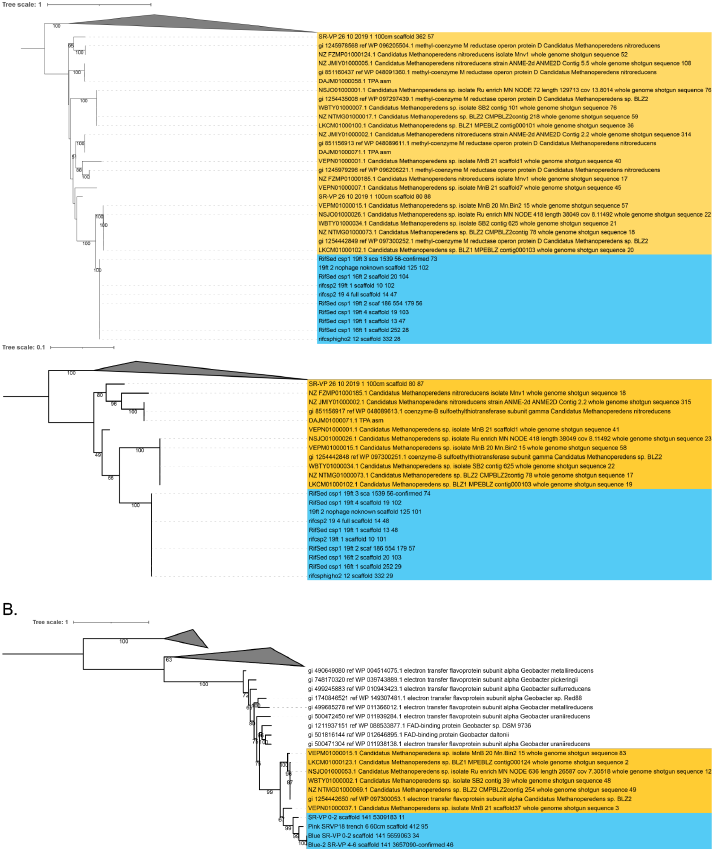
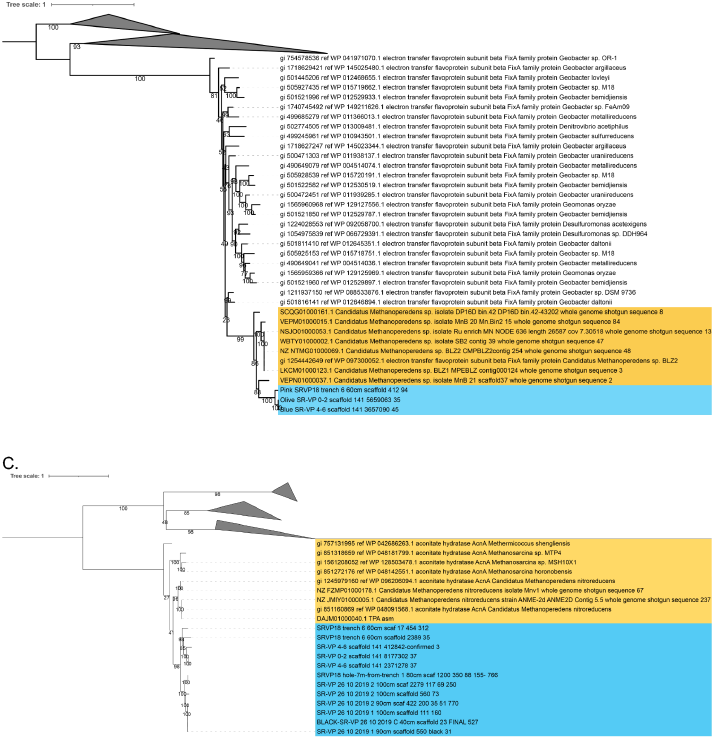
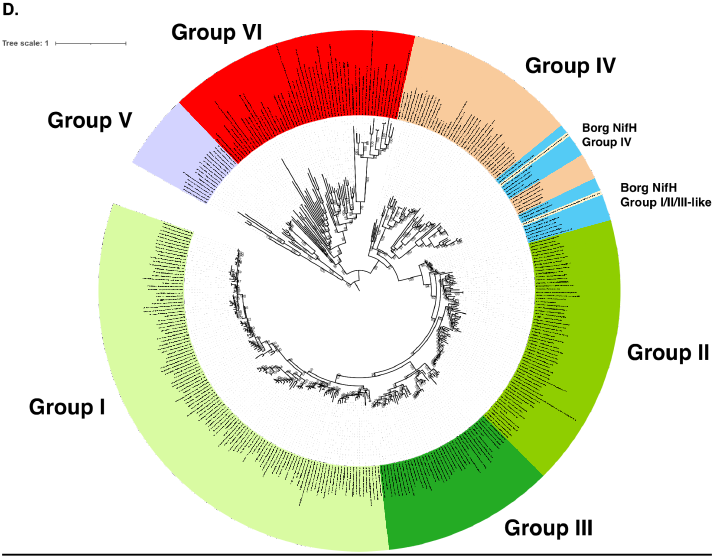
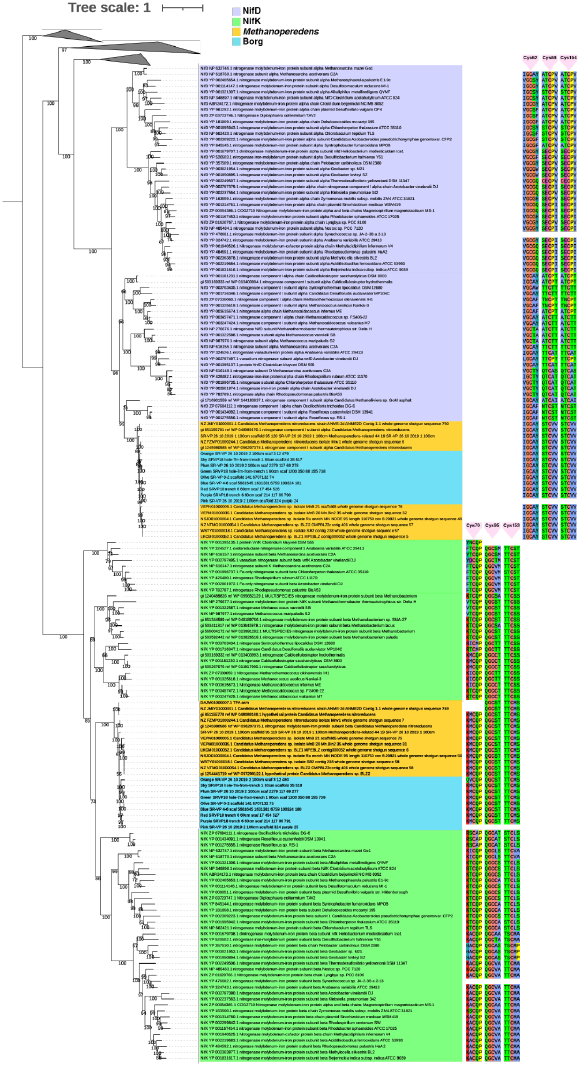
Phylogenetic trees for key Borg genes with functional predictions showing that the protein sequences (blue) cluster together and sibling to those from *Methanoperedens* (gold), with GC contents that approach those of *Methanoperedens*. (A) MCR_A, _B, _D, _G. (B) ETF Alpha, Beta. (C) Aconitase. (D) NifH and NifDK trees with alignments showing conservation of cysteine motifs that are involved in the attachment of the P-clusters.

### Supplementary Tables

**Table S1**: The complete, manually curated linear Borg genomes, partial genomes (bins) and bins only with short scaffolds that encode distinct Borg-like ribosomal protein L11 sequences. The Corona Mine outflow sampling site is located at 38°40’12” N, 122°32’09” W. The East River riverbed site is located at 38°55’24” N, 106°56’60” W and the Rifle CO site is located at 39°31’47” N, 107°46’20” W.

**Table S2**: The location, sequence, length and number of tandem direct repeats (3 or more units) for the four complete Borg genomes. In cases where two direct tandem repeat regions are adjacent and involve the same repeat sequence, the numbers of tandem direct repeats in each region are separated by a comma.

**Table S3**: Functional annotation overview.

**Table S4-S6:** Methanoperedens-specific, Borg-specific and protein families shared between Methanoperedens and Borgs.

**Table S7**: Gene name and description information for **Figure 3**. Sequences, specified by genome name and gene name, can be accessed via https://ggkbase.berkeley.edu/BMp/organisms

